# Spontaneous polarization and cell guidance on asymmetric nanotopography

**DOI:** 10.1101/2021.05.27.445842

**Authors:** Corey Herr, Benjamin Winkler, Falko Ziebert, Igor S. Aranson, John T. Fourkas, Wolfgang Losert

## Abstract

Asymmetric nanotopography with sub-cellular dimensions has recently been demonstrated to be able to provide a unidirectional bias in the migration of cells. The details of this guidance depend both on the type of cell studied and the design of the nanotopography. This behavior is not yet well understood, so there is a pressing need for a predictive description of cell migration on such nanotopography that captures both the initiation of migration and the manner in which cell migration evolves. Here, we employ a three-dimensional, physics-based model to study cell guidance on asymmetric nanosawteeth. In agreement with experimental data, our model predicts that asymmetric sawteeth lead both to spontaneous motion and changes in motion phenotypes. Our model demonstrates that asymmetric nanosawteeth induce a unidirectional bias in guidance direction that is dependent upon the actin polymerization rate and the sawtooth dimensions. Motivated by this model, an analysis of previously reported experimental data indicates that the degree of guidance by asymmetric nanosawteeth increases with the cell velocity.

---

**M**ost cells produce motion through the coupling of the actin cytoskeleton and the cell membrane to the surrounding substrate. *In vivo*, cell guidance is affected by a variety of external and internal stimuli, including the extracellular matrix (ECM) (1), surface rigidity(2), chemical gradients (3), and nanotopography. Local nanotopography has a substantial effect on cell behavior. For instance, it has been reported that nanotopographic substrates can lead to cell guidance and can influence cell morphology (4–7). Nanoridges can lead to the bidirectional contact guidance of cells, causing cells to speed up and elongate parallel to ridges (8). *In vivo* environments are often asymmetric (9, 10), in the form of the porous ECM. Such asymmetries are not often considered in modeling cell migration. Asymmetric nanotopographic substrates have recently been shown to lead to a unidirectional bias in cell migration, with a preferred guidance direction that depends on both the details of the nanotopgraphy (11) and cell line (12).

Physical modelling has been highly successful in describing cell migration, allowing for the creation of conceptually simple, whole-cell models with wide-reaching implications (13–20). Much of the previous modelling of cell migration has focused on cells on flat surfaces. However, the contact-guidance experiments discussed above suggest that the topography encountered in 3D, *in vivo* environments can have a profound effect on cell migration. We have previously developed a model that successfully captures some aspects of migration on nanotopographic substrates (21). Here we extend this model to rows of asymmetric nanosawteeth. The cell is modeled as a deformable boundary in which actin polymerizes at the front, forming lamellipodia and pushing the cell boundary outward. This extended model reproduces the key qualitative aspects of cell motility on asymmetric nanotopography.

Our work reveals that on asymmetric, nanoscale sawteeth, cells exhibit spontaneous onset of motion and directed guidance. For a cell in a deterministic model to move on a flat substrate, the cell must initially be polarized (13, 16, 19, 21). However, when the same cell model is implemented on asymmetric nanotopography, spontaneous cell polarization is readily observed. In agreement with previous experimental results (11, 12), our model exhibits biased unidirectional motion on asymmetric nanosawteeth, and exhibits a guidance direction that is dependent upon both the details of the nanotopography and actomyosin dynamics. Our model also reproduces qualitative cell-shape phenotypes observed experimentally, including cell elongation parallel and perpendicular to the guidance direction, depending upon the parameters of the nanotopography and the cells. The model also predicts that the degree of guidance by asymmetric sawteeth depends on cell velocity. This prediction is supported by a new analysis of previous experimental data.

## Results

### Phase-field Model

To explore cell migration on nanotopography, we extended a 3D phase-field model that was originally developed to model cells in confined environments (21). This model uses two dynamical variables to describe the state of the cell. The first, *ρ*(**r**, *t*), is a scalar phase-field that describes the cell boundary and assumes values between 0 and 1. The cell boundary lies at *ρ* = 1*/*2, with *ρ >* 1*/*2 inside the cell and *ρ <* 1*/*2 outside of the cell. The second is a 3D vector field **p**(**r**, *t*). The direction of **p** gives the orientation of the actin filaments and the magnitude of **p** gives the ordering of the filaments. As in the previous 3D phase-field model (21), we describe the substrate using two auxiliary fields: Φ(**r**) is a steric field that models the volume exclusion of the cell and the substrate and Ψ(**r**) restricts actin generation to occur close to the surface. The modeling framework is described by two coupled equations for *ρ* and **p**, as detailed in the Methods section. In the equations the unit of length is 1 *µ*m and the unit of time is 10 s.

Here we study the effect of three model parameters on guided cell migration and cell shape. The first, *β*, is the actin polymerization rate, which determines how fast actin polymerizes at the surface of the cell. The second, *θ*, determines the preferred angle of actin polymerization with respect to the local substrate geometry. When far away from the substrate, actin polymerizes normal to and away from the local cell membrane. Near the substrate, *θ* determines the component of actin polymerization normal to the substrate. For *θ* = 0, actin polymerizes tangentially to the local substrate, whereas for *θ* = 1 the normal and tangential polymerization rates are equal. The third parameter, *σ*, determines the rate of acto-myosin contraction. A larger value of *σ* represents more acto-myosin contraction in the cell. Collectively, these parameters can describe a wide variety of cell migration and shape phenotypes (21).

### Asymmetric nanotopography causes spontaneous polarization

We studied cell dynamics on an array of in-registry rows of asymmetric sawteeth, in which both the sawteeth and the spacings between rows were of subcellular dimensions (Fig. 1A). Such sawteeth have been shown experimentally to be able to bias cell guidance unidirectionally. Sun et al. (11) reported that different nanosawtooth heights lead to opposite guidance directions of *Dictyostelium discoideum*. Furthermore, Chen et al. (12) reported that the different cell lines were guided in opposite directions on nanosawteeth with the same height. For this study, the sawtooth height was varied between 0.75 *µ*m and 2 *µ*m and the sawtooth length was varied between 2.5 *µ*m and 6 *µ*m. We investigated how this nanotopography affects the onset of cell migration and the corresponding migration and shape phenotype evolution by placing a single cell on the substrate without initial polarization and observing the subsequent dynamics. The model produces two major cell-shape phenotypes, both of which are persistent. The crescent phenotype (Fig. 1B) is representative of more contractile cells, such as the M4 cell line that has been studied on asymmetric sawteeth (12). The wedge phenotype (Fig. 1C) is representative of cells on the lower end of the contractility spectrum, such as *D. discoideum*, which has also been studied on asymmetric sawteeth (11).

**Fig. 1.**
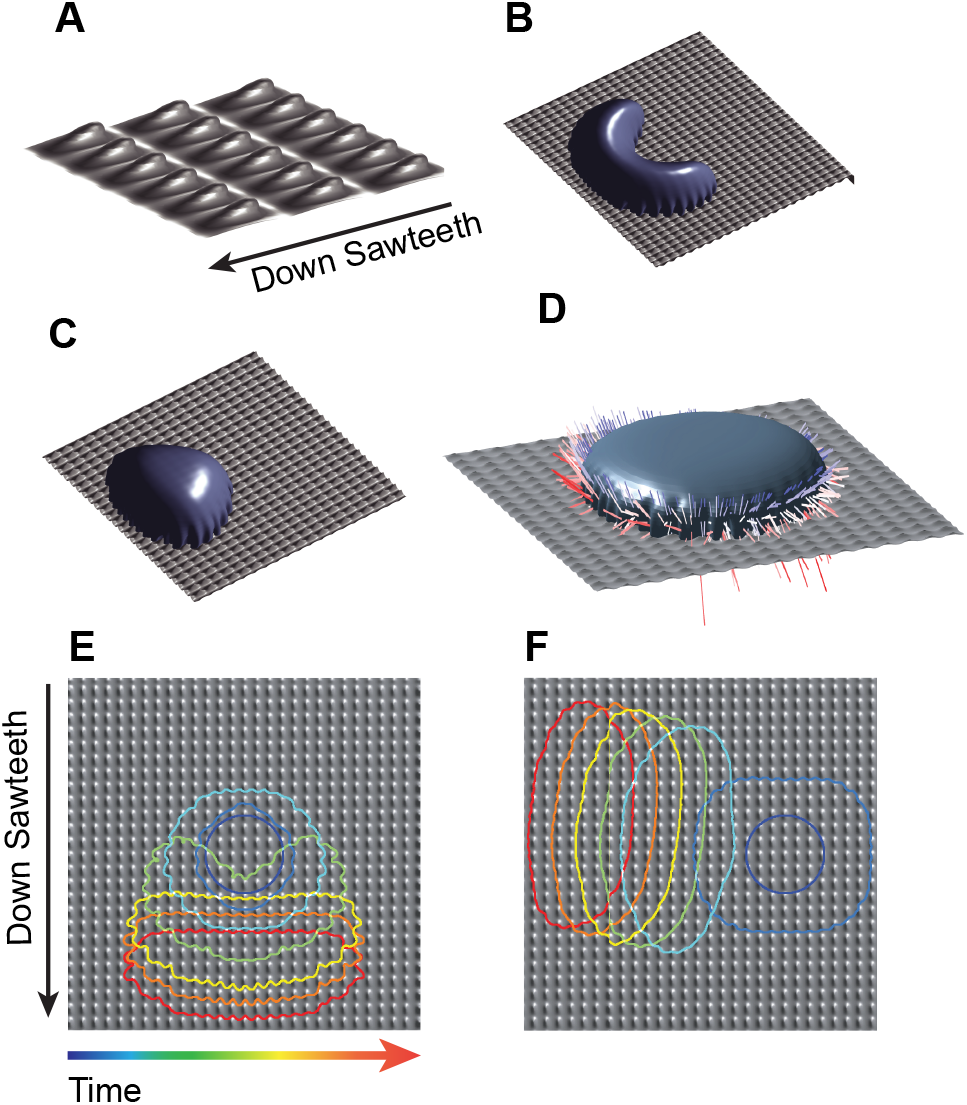
Asymmetric sawteeth and dominant phenotypes. (*A*) Zoomed-in schematic of the array of asymmetric sawteeth used in the phase-field model. Multiple cell-shape phenotypes are observed, including (*B*) crescents and (*C*) wedges.(*D*) Polarization vector field of a cell moving down the sawteeth. The polarization vector field indicates the directions of the actin polymerization and the membrane pushing force. The color denotes the strength of the polarization **p**, with blue being the weakest and red being the strongest. All polarization vectors point out of the cell. The asymmetric sawtooth nanotopography can cause the cell to polarize spontaneously (*E*) either down the sawteeth or (*F*) perpendicular to the sawteeth.

Spontaneous polarization is not observed in phase-field models of cells on flat substrates because the resting state is stable. Thus, the cells in this situation cannot migrate without inducing polarization externally (22) or via initial conditions (13, 23). However, the asymmetry of the substrates studied here allows cells to generate propulsion even in the absence of an initial polarization. Figure 1D shows the polarization vector field for a cell that is polarized spontaneously by sawteeth.

There are points of strong polarization underneath the cell that have a tangential component up the sawteeth (against the net direction of cell motion). However, there is strong enough polarization at the leading edge of the cell to propel the cell forward. This cell moves spontaneously with *θ* = 0. Therefore, the impetus for spontaneous motion on sawteeth is actin polymerization tangential to the substrate. In this model, a cell can only move spontaneously down the sawteeth (Fig. 1E) or at an angle to the sawteeth (Fig. 1F).

### The direction of cell guidance is influenced by actin polymerization

In experiments with cells on asymmetric nanosawteeth, the guidance direction has been observed to depend on the sawtooth height and length (11), as well as on the cell type (12). We next explore why different types of cells can be guided in different directions on the same sawtooth pattern. When inducing an initial polarization up the sawteeth, we observed three possible outcomes in our model: the cell moves up the sawteeth persistently (Fig. 2A), the cell initially moves up the sawteeth and then turns around to move down the sawteeth (Fig. 2B), or the cell stops. We induce the polarization up the sawteeth to counter the preferred spontaneous polarization down the sawteeth. Trajectories for these three representative behaviors are shown in Fig. 2C.

**Fig. 2.**
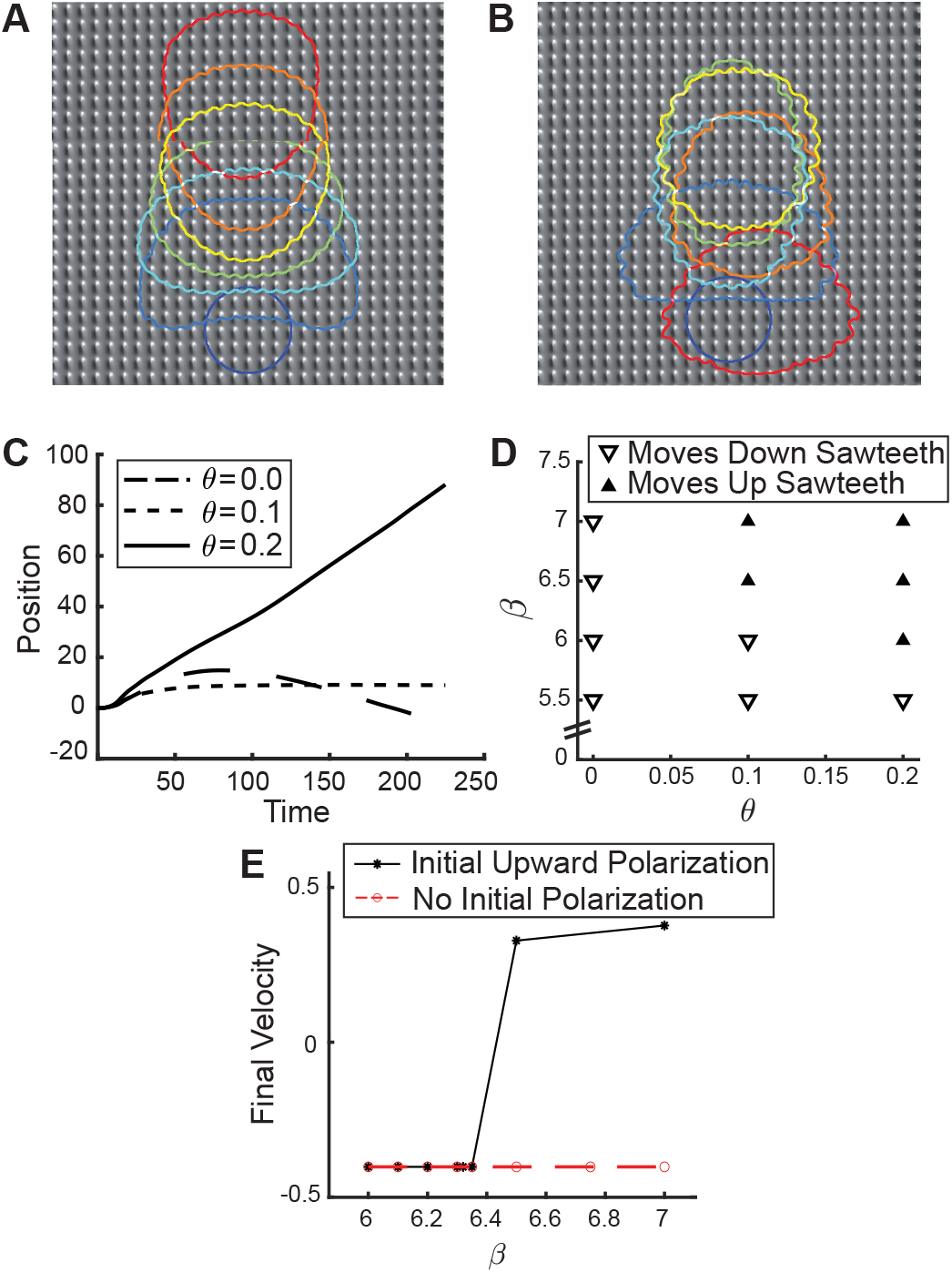
Reversal of direction and evolution of shape phenotype. (*A*) A cell guided unidirectionally up the sawteeth following initial polarization up the sawteeth. (*B*) Decreasing the actin polymerization rate leads to a trajectory in which the cell initially moves up the sawteeth but then turns around. (*C*) Trajectories representing the three most common outcomes when a cell on the sawteeth is polarized up the sawteeth. (*D*) A phase diagram showing how the guidance direction depends on the actin polymerization rate *β* and the angle *θ* of actin polymerization relative to the substrate. The cell moves up the sawteeth when both *β* and *θ* are in the larger range of the values examined here. (*E*) The asymptotic guidance velocity exhibits hysteresis for *θ* = 0.1. For cells with an initial polarization, the transition from moving up the sawteeth to moving down the sawteeth is sudden. For unpolarized cells, guidance is only observed down the sawteeth.

To probe the selective guidance direction systematically, we created a phase diagram (Fig. 2D) in which we varied *β* and *θ* while keeping the sawtooth parameters the same. In all cases the cell had an initial upward bias. After 250 seconds, we measured the center-of-mass velocity. This velocity was used to determine the guidance direction shown in Fig. 2D. The data in this figure indicate that as the actin-polymerization rate increases, the guidance direction changes from down the sawteeth to up the sawteeth.

We studied the transition in guidance direction by changing *β* at a fixed *θ* of 0.1. Figure 2E shows that for a cell with an initial polarization, there is a sharp transition of guidance direction at *β* = 6.5. Similar transitions occur at other values of *θ*. Below *β* = 6.5, the cell always has the same final (asymptotic) velocity. This guidance velocity is determined by the sawtooth properties, rather than by the model parameters. Additionally, we observe hysteresis in the guidance direction. If the cell is not polarized initially, the cell can only be guided down the sawteeth. This observation indicates that the preferred guidance direction is down the sawteeth. Finally, a cell that reverses direction does not move solely parallel to the sawteeth. We find instead that a cell can turn to move at an oblique angle to the sawtooth orientation.

### The substrate parameters can be used to control the cell shape and the guidance direction independently

Our model exhibits a range of stable shape phenotypes that were studied in relation to the guidance direction. To control shape, we modified the contractility of the cell, *σ*. Contractile forces control the shape of the cell (24), and are linked closely to cell motility (25). Here, we investigate the effect that contractility has on the direction of guidance. Figure 3A depicts a phase diagram of guidance direction as a function of *σ* and *β* (*θ* = 0.1). Cells with higher contractility *σ* and higher actin polymerization rate *β* move up the sawteeth, counter to the preferred direction of spontaneous motion.

**Fig. 3.**
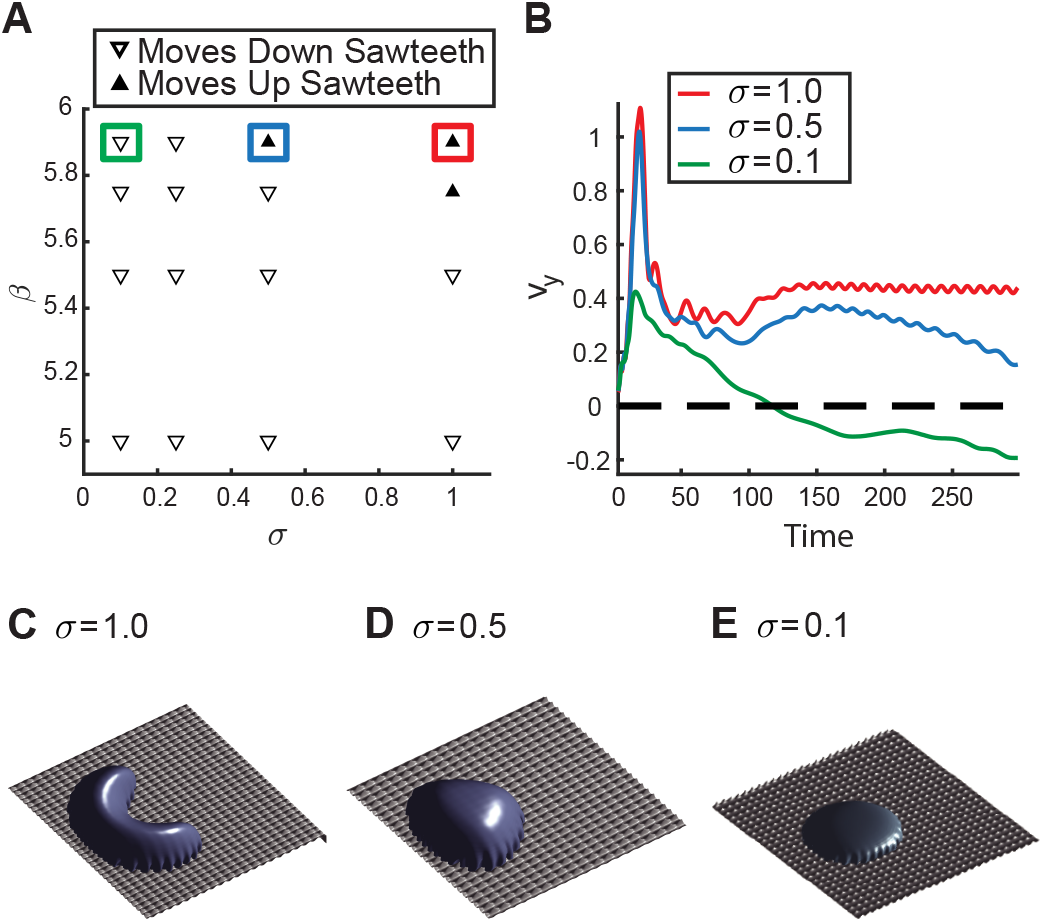
The mode parameters can be used to control shape and guidance independently. *A*) A phase diagram showing the guidance direction as a function of the contractility *σ* and the actin polymerization rate *β* (*θ* = 0.1). Increasing the contractility leads to a reversal in the guidance direction. (*B*) Cell migration behavior for different values of *β* and *σ*. For *σ* = 1, the cell’s initial velocity decays rapidly to a more stable final velocity. For *σ* = 0.5, the cell’s initial velocity decay mirrors that for *σ* = 1, before decaying further at longer times. For *σ* = 0.1, the cell reverses direction at *t* ≈ 100 to move down the sawteeth. 3D visualizations of the stable cell-shape phenotype for (*C*) *σ* = 1, (*D*) *σ* = 0.5, and (*E*) *σ* = 0.1. As the contractility *σ* decreases, the cell becomes more oval shaped.

By examining the dynamics at individual points in the phase diagram (Fig. 3B), we can separate the velocity trajectories into two distinct regions: the first peak and dip in velocity due to the initial polarization of the cell, and the ensuing long-term behavior. The initial peak and dip occur for all values of *σ*, followed by a positive rebound in velocity that occurs at *t* ≈ 150. The major differences among the trajectories are in the long-time behavior. For *σ* = 1 there is a small negative acceleration, but the velocity up the sawteeth is persistent. The velocity is oscillatory because the cell slows down as it crawls up the sawteeth at its leading edge. The persistent velocity and the strong contractility of the cell results in elongation perpendicular to the direction of motion and leads to a persistent crescent shape (Fig. 3C). For *σ* = 0.5, the long-time positive velocity is not persistent. During the period of the simulation the cell moves up the sawteeth, but with a steadily decreasing velocity at long times. The contractility is high, so the cell is again elongated perpendicular to the direction of motion (Fig. 3D). Finally, for *σ* = 0.1, the long-time velocity is negative. A short period of positive acceleration at *t* ≈ 175 is observed but does not lead to a second change of direction. The cell in this case has an oval shape that is elongated parallel to the direction of motion (Fig. 3E).

### Comparison to experimental cell shapes and motion

Next we reanalyze previous experimental data on *D. discoideum* migration on asymmetric sawteeth to capture additional details of cell motion. We find that the deterministic phase-field model can reproduce both the observed cell shapes and the preferred guidance direction. Figures 4A-B show that *D. discoideum* can be elongated both parallel and perpendicular to the sawteeth, in analogy to the shape phenotypes seen in the model. *D. discoideum* cells moving along sawteeth tend to elongate in the direction of motion, in accordance with the less contractile cells shown in Fig. 3. Additionally, in a study of different cell lines migrating on sawteeth by Chen et al. (12), it was observed that M4 cells, a cancerous mutant of MCF10A epithelial cells, exhibit elongation perpendicular to the direction of motion. This behavior agrees with the observation that cancerous cells are generally more contractile than their progenitor cells (25). Although asymmetric sawteeth create a unidirectional bias of motion, *D. discoideum* is capable of moving down the sawteeth (Fig. 4C), up the sawteeth (Fig. 4D), as well as at arbitrary angles relative to the sawteeth.

**Fig. 4.**
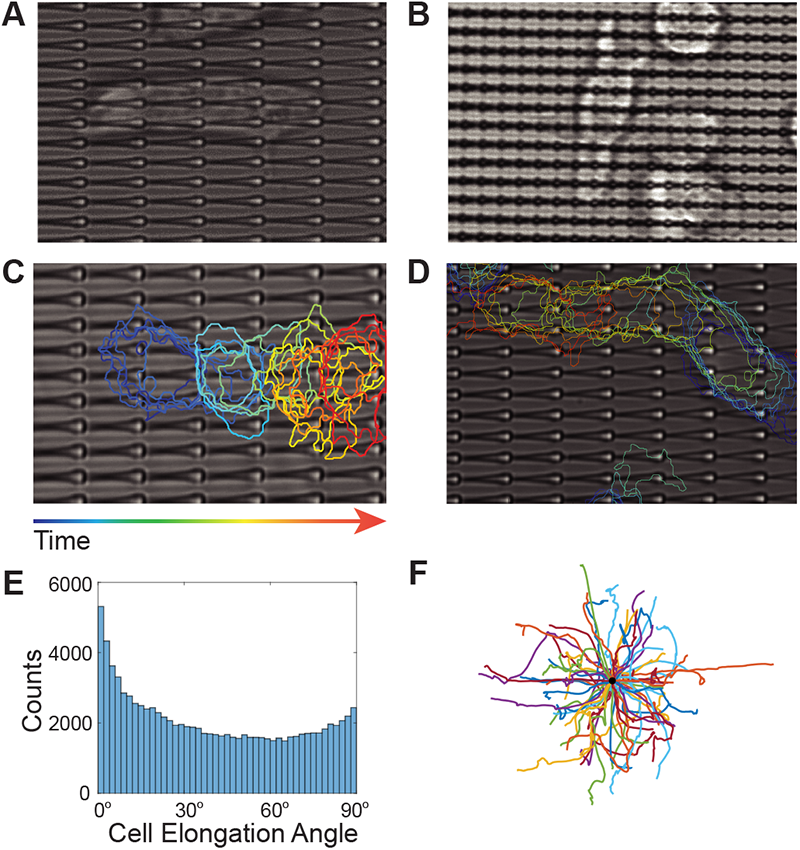
Experimental data support the predictions of the model. *D. discoideum* cells on asymmetric sawteeth are observed to elongate both (*A*) parallel and (*B*) perpendicular to the sawtooth direction during cell migration. Motion of *D. discoideum* (*C*) down and (*D*) up the sawteeth. (*E*) The full distribution of elongation angles of *D. discoideum* on asymmetric sawteeth. The distribution is peaked around the parallel and perpendicular directions, as seen in the model. However, cells can exhibit other orientations as well. (*F*) The full distribution of directions of motion of *D. discoideum* on asymmetric sawteeth of heights 1 *µ*m to 2.4 *µ*m. The model can also produce the full range of directions of motion.

A variety of cell elongation angles with respect to the sawtooth orientation are observed experimentally, as seen in Figure 4E. The distribution of elongation angles has two local maxima at 0° (parallel to sawteeth) and 90° (perpendicular to sawteeth). Fig. 4F shows a combined spider plot of cell trajectories on sawteeth of height 1 *µ*m to 2.4 *µ*m. By changing the direction of initial cell polarization, a model trajectory can be produced that travels in any direction, and therefore the model can reproduce the arbitrary elongation angles that are present experimentally.

The simulated cell trajectories in the continuum model are deterministic, and so cannot reproduce the stochasticity of experimental trajectories. Nevertheless, we observe that the direction of stable simulation trajectories corresponds to the preferential guidance direction of cells on asymmetric saw-teeth. To determine the effect of sawtooth dimensions on cell guidance direction we compared the model to experiments of cells on sawteeth of three different heights: 1 *µ*m, 1.8 *µ*m, and 2.4 *µ*m. The model predicts that only cells with high actin polymerization rates will move up the higher sawteeth (Fig. 5A). To determine the guidance direction of cells in experiments, we measured the average velocity of the *D. discoideum* cells as a function of direction relative to the sawteeth, with the angle 0° defined as moving up the sawteeth (Fig. 5B-D). For the 1 *µ*m-high sawteeth, the average velocity peaks at 0° (Fig. 5B). This observation is consistent with the phase diagram (Fig. 5A), in which cells with a higher actin polymerization rate move up the sawteeth. For the 1.8 *µ*m-high and 2.4 *µ*m-high sawteeth the average velocity is peaked at 180°. This result indicates that the *D. discoideum* cells do not reach the threshold velocity necessary to move up these sawteeth. Our observations suggest that asymmetric sawteeth can be used to guide cells with high velocities selectively, and that the velocity cutoff can be modified by changing the sawtooth height.

**Fig. 5.**
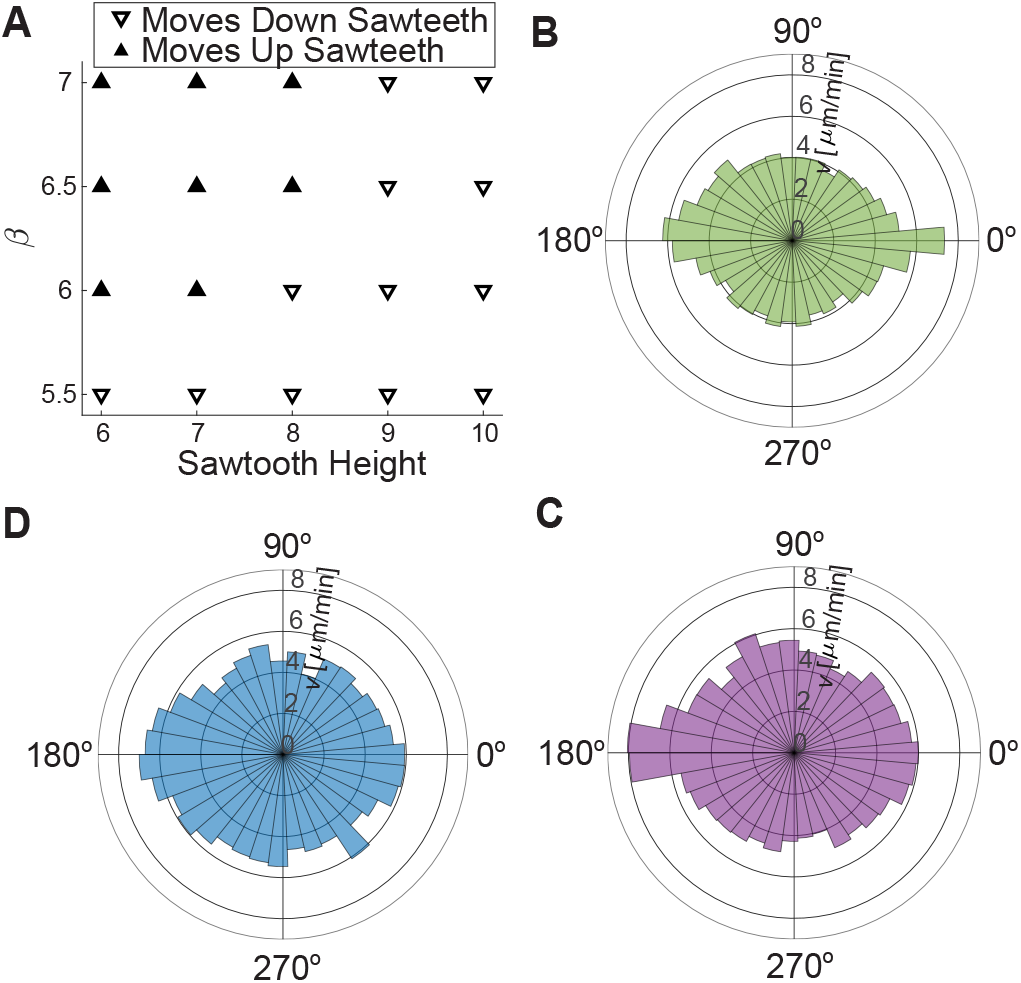
Asymmetric sawteeth preferentially guide cells with high velocities. (*A*) A phase diagram demonstrating dependence of guidance direction on *β* and sawtooth height in the phase-field simulations. The average velocity distributions derived from experimental data for *D. discoideum* on asymmetric sawteeth of three different heights: (*B*) 1 *µ*m, (*C*) 1.8 *µ*m, and (*D*) 2.4 *µ*m. For the 1*µ*m sawteeth, the distribution is peaked at 0, indicating that the faster cells are guided preferentially up the sawteeth. This observation is consistent with the phase diagram in (*A*), in which cells with higher actin polymerization rates are guided up the sawteeth. For sawtooth heights of 1.8 *µ*m and 2.4 *µ*m, the velocity distribution is peaked at 180*°*, indicating that the faster cells move preferentially down the sawteeth.

## Discussion

### A toy model for cell migration on nanotopography

Important insights into cell migration can be obtained from a simple model describing a cell as a material point with a velocity threshold for the onset of motion. If there is no topography on the substrate, the equations of motion can be cast in the dimensionless form:

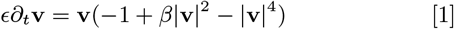

Here *E "* 1 is the relaxation parameter and *β* is the driving parameter related to actin polymerization. For *β >* 2, equation (1) possesses 3 stable fixed points: *V*_0_ = 0, and 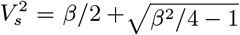 Correspondingly, there are two unstable fixed points 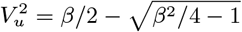.

In the presence of asymmetric nanotopography, the equation of motion assumes the form

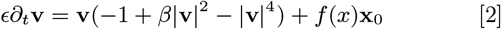

Here **x**_0_ is the unit vector in *x*-direction. In the case of asymmetric sawteeth, *f* (*x*) can be written in the form

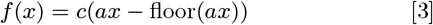

Here *a* is the spatial frequency (spatial period *L* = 1*/a*) and *c* is the magnitude of the sawtooth modulation. Note that the sawteeth provide a positive bias, because *f* (*x*) *>* 0.

Despite their simplicity, Eqs. (2) and (3) capture many salient features exhibited by the fully 3D, continuum phase-field model as a function of the driving parameter *β* (Fig. 6). In particular, for pure one-dimensional motion (i.e., in the *x*-direction), for a large enough value of *β* the model exhibits two stable solutions, with motion up or down the sawteeth (Fig. 6A). For smaller values of *β*, the model only shows spontaneous motion down the sawteeth. For even smaller values of *β*, no motion is possible, all in agreement with the phase-field model. The two-dimensional version of Eqs. (2) and (3) yields an intriguing prediction that motion up sawteeth is always unstable (Fig. 6B). A cell that is initially polarized in the direction up the sawteeth will, after some time, make a U-turn and begin to move down the sawteeth. However, as *β* increases the reversal time diverges (Fig. 6C). Thus, the model indicates that the complex migration behavior observed both in the phase-field simulations and in the experiments is a synergistic effect of the sawtooth substrate and the cell’s intrinsic threshold for motion.

**Fig. 6.**
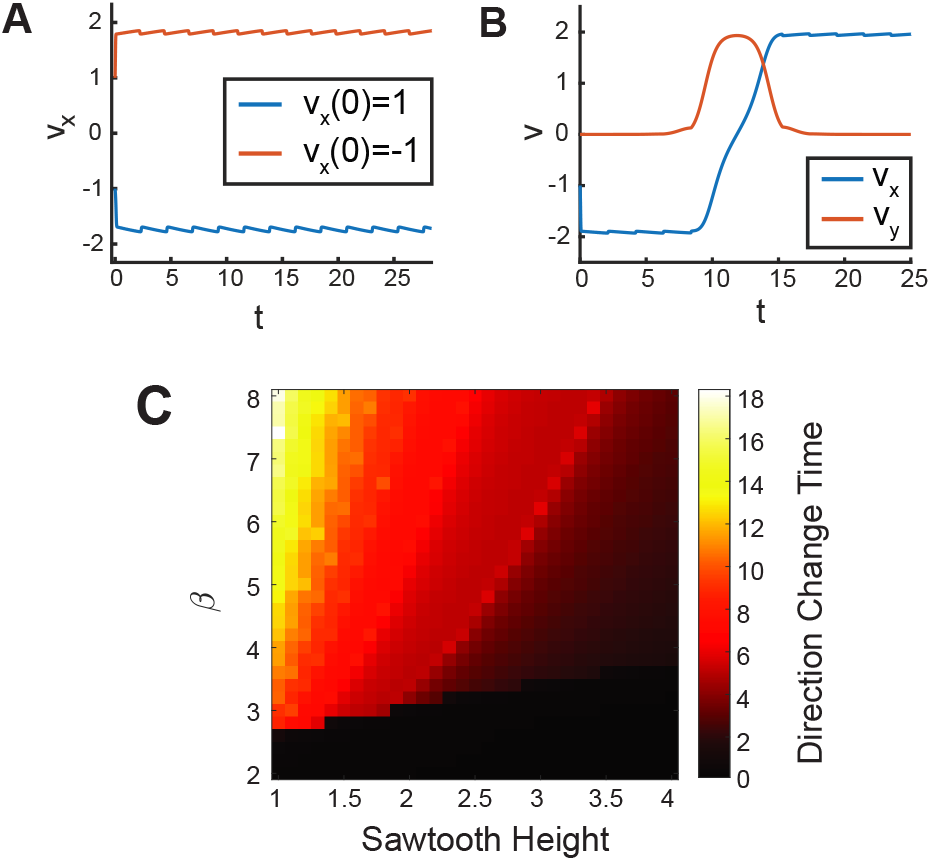
Toy model capturing important features of full 3D phase-field model. (*A*) Trajectories up and down the sawteeth in the simplified model, Eq. (2), in 1D case. (*B*) A cell that initially moves up sawteeth in the 2D model, Eq. (2) performs a U-turn at *t* ≈ 10. The initial trajectory is unstable. (*C*) Heat map of the time for cells with initial negative velocity to change direction in the 2D model. For higher values of *β* in this model, the velocity up the sawteeth persists for a longer time. For higher values of *c* (height of sawtooth pattern) in this model, the velocity up the sawteeth persists for a shorter time

## Conclusions and outlook

We have demonstrated that a 3D phase-field model enables a rigorous investigation of cell motility on asymmetric nanotopography. The phase-field model captures the spontaneous onset of motion and the experimentally observed range of shapes and motions of cells that sense asymmetric guidance cues that are an order of magnitude smaller than the scale of the cell.

Cell-migration experiments on asymmetric sawteeth have demonstrated that the guidance direction is affected by the sawtooth dimensions (11) and the specific cell line used (12). The phase-field model reproduces the key phenomena observed in these experiments, including spontaneous polarization, unidirectional guidance, and elongation phenotypes. By tuning model parameters, we can change the guidance direction and cell shape in accordance with the behavior observed for different cell lines. The correspondence between the model and experiments yields insights into the relative importance of actin polymerization, contractility, or other cell properties in promoting cell-type-dependent guidance behavior.

On flat surfaces, the phase-field models require an initial polarization (13, 21) or a strongly randomized internal actin density (26) to observe directed cell motion. In contrast, asymmetric sawteeth provide guidance cues that cause the cell to move spontaneously. Consequently, surface asymmetries may be one factor that initiates directional migration *in vivo* and in natural environments.

Our model shows that the actin polymerization rate is a primary driver of the cell guidance direction on asymmetric sawteeth. There is a transition in guidance direction from down the sawteeth to up the sawteeth at a critical value of the actin polymerization rate. This critical value is dependent on other model parameters as well, such as the sawtooth height. The actin polymerization rate is directly related to the cell velocity, so faster cells should tend to move up the sawteeth. In agreement with the predictions of our model, experimental data for cell guidance on asymmetric sawteeth show velocity-dependent selective guidance. Additionally, the model predicts that increasing the sawtooth height should increase the velocity threshold for cells to move up the sawteeth. We verified this prediction by showing that faster *D. discoideum* cells are preferentially guided up asymmetric sawteeth that are 1 *µ*m high. However, on asymmetric sawteeth with heights of 1.8 *µ*m and 2.4 *µ*m, faster cells are still guided down the sawteeth.

In our model, contractility affects both the shape and the guidance direction of a cell. Larger values of contractility are associated with the cell moving up the sawteeth. Thus, more contractile cells are not guided in the more typical direction on asymmetric sawteeth. Studies have linked higher contractility to invasiveness in cancer cells (25), and this connection may be related to the change in guidance direction that we.

The simple physical model presented here can explain many features of cell motility on asymmetric nanotopography. Because the phase-field model is highly modular, it can be extended further to obtain a fuller picture of cellular dynamics on asymmetric nanotopography. An even more realistic model could include the modulation of specific or non-specific adhesive effects (e.g., focal adhesions). Additionally, the actin dynamics can be modified to include the effect of regulatory proteins via an excitable network model such as local excitation global inhibition biased excitable network (27) or linked excitable networks (28).

Although our approach captures many important aspects of subcellular environmental asymmetries on cell migration, *in vivo* the dynamics can be affected by many other factors. It is known that the cells can remodel, and even degrade, the local topography, especially in the context of some invasive cancers (29, 30). Many cancer cells form invadopodia, i.e., actin-rich protrusions of the plasma membrane that are associated with degradation of the extracellular matrix. Incorporation of such phenomena into a phase-field model can shed light on cancer invasiveness and metastasis and is highly desirable. However, a 3D phase-field model on deformable and degradable substrates is a formidable computational task that we leave to future study.

## Materials and Methods

### Phase-field equations

The following model was used to describe the cell migration in 3D:

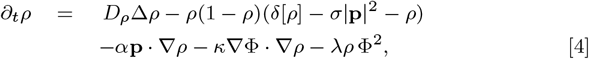

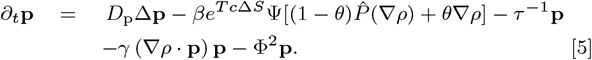

Here, the phase-field *ρ* describes the cell shape, **p** characterizes the actin orientation, Φ(**r**) models a steric field moderating the cell’s interaction with the surface, and Ψ(**r**) restricts actin generation close to the surface. The entire motility mechanism is captured by Eqs. (4), (5). Actin polymerizes near the cell boundary and produces a protrusion force via the polymerization ratchet mechanism(31). As the cell moves forward, the actin polarization at the back of the cell degrades based on a model of acto-myosin contraction (13). The cell volume is conserved as a fixed point through the term *δ*[*ρ*]. This model of cell motility has been shown to reproduce the lamellipodium-based motion of keratocytes on flat surfaces (13). We use a modified version of the model that accounts for cell surface tension (32). Surface tension is introduced using the exponential term *e*^-*T c*Δ*S*^, where *T* is the surface tension strength, *c* is the curvature, and Δ*S* is the difference between surface area at a given time step and a reference surface area. This term makes the actin polymerization rate lower at points of high curvature, thereby preventing the cell from tearing apart. We introduced this term by necessity based on the high curvature of the nanotopography studied here.

### Numerical Method

The model used here is an extension of a previously published method (21). Equations (4) and (5) were solved on a 100 *µ*m × 100 *µ*m × 25 *µ*m rectangular grid (corresponding to 576 × 576 × 128 mesh points) in the domain using a split operator method on a periodic domain. The algorithm was implemented on graphical processing units (GPUs) using the NVIDIA CUDA programming language. The diffusion terms were calculated in Fourier space, and other operators were calculated using finite-difference methods. The curvature *c* in Eq. (4) was calculated using 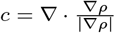. Unless otherwise mentioned, the model parameters used were those given in Table 1. The substrate with asymmetric sawteeth was implemented via prescribing the static fields Φ and Ψ. The substrate studied for this paper was a sawtooth of length 4.2 *µ*m and a default height 1.4 *µ*m; other heights were used as discussed above.

**Table 1.** **Model Parameters**

| Parameter | Value | Description |
| --- | --- | --- |
| $D_\rho$ | 1 | Diffusion constant for $\rho$ |
| $D_p$ | 0.2 | Diffusion constant for $\mathbf{p}$ |
| $\alpha$ | 1.5 | Actin pushing strength |
| $T$ | 25 | Surface tension strength |
| $\gamma$ | 1 | Front-back asymmetry |
| $\tau^{-1}$ | .1 | Actin degradation rate |
| $\kappa$ | 2 | Surface adhesion strength |
| $\theta$ | 0.1 | Actin polymerization angle |

### Cell Tracking

Cell tracking was performed on bright-field movies of *D. discoideum* moving on asymmetric sawteeth reported in a previous paper (11). First, the threshold of the fast Fourier transform (FFT) of the images was used to remove the bright spikes associated with the periodic sawteeth. A low-pass Hamming filter was applied to the FFT. An inverse FFT was then applied to recover an image with sawteeth removed. Next, the image contrast was changed so that 1% of the brightest and darkest parts of the image were saturated. A morphological top-hat filter was applied to remove high frequency noise. Finally, the image was binarized using the Otsu thresholding method (33). The cells were tracked using the MATLAB *regionprops* function to find elongation and centroid location. The cell trajectories were found using a particle tracking algorithm fed the centroids of the cell locations.

## Notes

### Competing Interest Statement

The authors have declared no competing interest.

